# *ninna nanna* links circadian and homeostatic sleep drive in *Drosophila*

**DOI:** 10.1101/2024.05.10.593616

**Authors:** Anne Petzold, Giorgio F. Gilestro

## Abstract

Sleep is under control of two processes – circadian and homeostatic regulation – but little is known about how these two processes integrate. Here, we identify a *Drosophila* gene, *ninna nanna*, encoding two alternatively spliced isoforms: Ninna and Nanna. Both proteins encode aldo-keto reductases and are expressed in different, yet interconnected neurons. One isoform, *ninna*, encodes an aldo-keto reductase with predicted affinity for NADP(H) and is expressed in key circadian pacemaker neurons, the s-LN_v_s. The second isoform, *nanna*, encodes an aldo-keto reductase with predicted affinity for NAD(H) and is expressed in ICLI neurons, a pair of wake-promoting peptidergic neurons whose inhibition relieves sleep pressure. Ninna and nanna neurons interconnect to integrate a binary sleep sensing circuit in which *ninna* receives circadian information and *nanna* encodes homeostatic sleep pressure. *ninna nanna* defines an archetypal circuit for the integration of circadian and homeostatic sleep drive and reinforces the hypothesized link between aldo-keto reductases and sleep regulation.

## Introduction

Across the animal kingdom, the amount and timing of sleep are regulated by an internal “somnostat”, operated by the concerted activity of at least two processes: a circadian process (process C) and a homeostatic process (process S) (1, 2). In the classical formalisation of the so-called two-process model, the homeostatic process regulates sleep pressure as a function of awake time – the longer the time spent awake, the stronger the sleep pressure – while the circadian process is responsible for setting the threshold for sleep (3). At any given time, the propensity to sleep can therefore be defined as the mathematical difference between S and C (4). Besides providing a framework for describing sleep pressure, the formalisation of the two-process model has proven to be instrumental to define many corollary aspects, such as the electrophysiological correlates of sleep (5), and some cognitive correlates of waking (6). However, while the two-process model provides a robust theoretical description of how the circadian and homeostatic controllers cooperate, little is known about the biological underpinnings of the somnostat. How do C and S physically integrate to regulate when and how an animal should sleep?

While the cellular and molecular regulation of circadian rhythms has largely been identified (7–9), the biological mechanisms that ensure sleep homeostasis are just beginning to be understood (10). In mammals, the *flip-flop* switch model (11) provides a widely accepted, yet incomplete, sketch of a circuit regulating sleep. Briefly, the circadian rhythm of sleep and wakefulness is under control of the neurons of the suprachiasmatic nucleus (SCN) in the hypothalamus, impinging on the neurons of the ventrolateral preoptic nucleus (VLPO), locus coeruleus (LC), as well as orexin/hypocretin neurons (HONs) of the lateral hypothalamus, to ultimately flip the binary state between sleep and wakefulness. The *flip-flop* model does not involve a neuronal circuit specifically encapsulating homeostatic sleep pressure, given that none has been identified. Instead, adenosine – a key by-product of neuronal metabolism primarily formed from the breakdown of adenosine triphosphate (ATP) – has been proposed as a widespread molecular regulator of sleep drive (12–14) that ultimately modulates the activity of VLPO, LC and HONs through adenosine receptors expressed on the cell surfaces (15, 16).

A conceptually similar model is emerging for *Drosophila melanogaster*. In flies, sleep timing, duration, and structure are also controlled by the anatomical and functional integration between circadian pacemaker neurons and a number of homeostatic sensing neurons (14, 17–24). The circadian gears of this mechanism are well understood due to decades of pioneering research in this field, but the other side of this coin – the molecular and cellular mechanisms that encode sleep pressure – remain poorly characterised.

An emerging hypothesis suggests that flies, akin to humans, might regulate their somnostat by integrating circadian pace-making neurons with intracellular metabolic sensors, but instead of relying on adenosine – like mammals – flies are proposed to utilize nicotinamide adenine dinucleotide phosphate (NADPH). First evidence that NADPH may be playing a crucial role came from the identification of *shaker* (25), the very first gene identified as a regulator of sleep in *Drosophila*, encoding for the *α* subunit of a voltage-gated potassium (K_v_) ion channel. Modulators of the Shaker channel have subsequently been confirmed as sleep-regulating proteins in the past two decades. These include the GPI-anchored protein Sleepless/Quiver (26–28), and the K_v_ channel *β* subunit Hyperkinetic (Hk) (29–31). Besides being a structural component of the channel itself (32), the latter plays an important regulatory role through its poorly understood enzymatic aldo-keto reductase (AKR) domain (32, 33). Aldo-keto reductases (AKRs) are a group of structurally related monomeric oxidoreductases (34) catalysing the conversion of carbonyl substrates (such as aldoses and ketoses) to alcohols using NADH or NADPH as cofactors (35). Rather than than actively engaging in enzymatic activity, however, the AKR domain of K_v_ channel *β* subunits are believed to be acting as redox sensors employed by the channel to read the ratios of NADH/NAD^+^ and NADPH/NADP^+^ (36, 37). The NADPH/NADP^+^ ratio in the central nervous system of *Drosophila* increases with sleep pressure (38) and the K_v_ *β* subunit Hk was proposed to act as a sensor for cytoplasmic levels of NAD(P)H/NAD(P)^+^ (39) to slow the inactivation of Shaker and increase neural activity facilitating sleep (39). Interestingly, while the removal of the Shaker potassium channel severely affects sleep (25), the removal of Hyperkinetic leads to a considerably milder sleep loss (29, 39), suggesting further players may be involved. Here, we show that another AKR gene, *ninna nanna*, tracks sleep need and potentially modulates sleep through an output node of the circadian pacemaker network in *Drosophila. ninna nanna* is a unique gene that expresses two alternatively spliced isoforms in different, yet complementary neurons, interconnecting into a novel circuit. Our findings suggest a general role for AKR genes as molecular sensors of sleep homeostasis.

## Results

### ninna nanna is a novel AKR gene regulating sleep in Drosophila

To identify AKRs that were expressed in neural circuits of the fly brain, we screened *Drosophila* AKR transcripts for their expression level in the adult fly brain using quantitative real-time PCR (qRT-PCR) of brain or body extracts of adult male flies. Only two AKRs are overexpressed in the brain of the adult animal: *hk* and the two alternatively spliced isoforms of the previously uncharacterised *CG10638* gene (Fig. 1A,B). The absolute expression level of these transcripts in the head is relatively low, indicating restricted expression patterns (Fig. S1). The single cell transcriptome atlas indicates expression in 5240 out of 56902 neurons (40).

**Figure 1.**
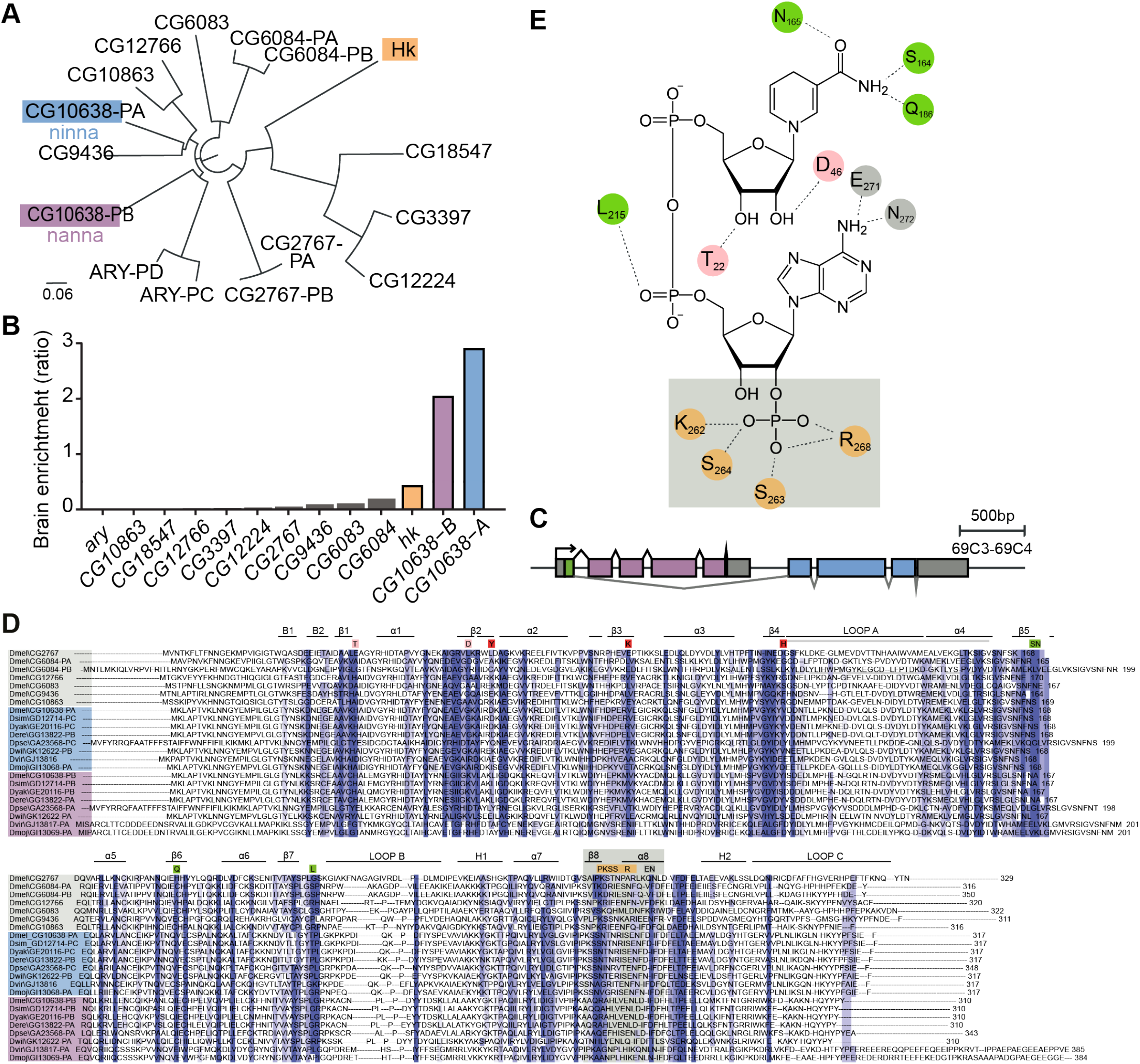
Brain-specific enrichment of specific AKR genes in *Drosophila melanogaster*. **A**, Phylogenetic tree of the 12 predicted AKR genes in *Drosophila melanogaster* including Hk (orange), CG10638 isoform A (blue) and CG10638 isoform B (purple). CG10638, CG2767 and CG6084 give rise to two alternatively spliced isoforms each. The Hk gene also undergoes alternative splicing to give rise to several, largely similar isoforms (not shown). Multiple sequence alignment (MSA) was generated using MUSCLE Clustal 2.1. **B**, Adult enrichment of AKRs in protocerebrum relative to the rest of the body assessed using quantitative real-time PCR (qRT-PCR), n = 3–4 biological replicates per tissue consisting of 12 brains or 5 headless bodies per sample. **C**, Schema of the *ninna nanna* locus. Boxes indicate exons; green labels the shared coding sequence; blue and purple label ninna and nanna specific coding sequences, respectively; grey labels non-coding sequences in exons. **D**, Alignment of the *Drosophila melanogaster* AKR proteins (grey background) and the conserved Ninna and Nanna proteins across eight Drosophilae species. The black lines above the sequences highlight the putative conserved tertiary structure. The residues highlighted are the ones critical for interaction with the co-factors. Red residues are the ones constituting the active site. **E**, Conserved amino acid important for AKR interaction with the NADP(H) co-factors. Three out of the four amino acids interacting with the phosphate group of NADP(H) are mutated in all Nanna proteins, i.e. orange residues located in *β*-sheet 8.

In *Drosophila melanogaster*, the *CG10638* gene — hereby referred to as *ninna nanna* after the Italian term for lullaby — is located on the third chromosome and is composed of 10 exons, splicing into two alternative isoforms: CG10638-RA (Ninna, blue in all figures) and CG10638-RB (Nanna, purple in all figures) (Fig. 1C). Although the *ninna* and *nanna* transcripts arise almost entirely from different exons, they both encode two predicted AKR enzymes with an identity of 57 % at the amino acid level (Ninna 317 aa; Nanna 310 aa; shared exon encodes for 24 aa; Fig. 1D). Interestingly, Nanna carries several conserved substitutions in three key residues that are predicted to interact with the phosphate group of NAD (S263A, S264A and R268H – orange residues in Fig. 1E), conferring a predicted preference for NAD(H) over NAD(P)H (41–44).

To test whether *ninna nanna* is involved in sleep regulation, we tested flies carrying a P-element insertion in the common exon (EP21723, green arrow in Fig. 2A). Mutant flies showed a reduced expression of both transcripts (Fig. 2B) and reduced sleep levels (Fig. 2C,D) but an intact circadian rhythm (Fig. 2E,F).

**Figure 2.**
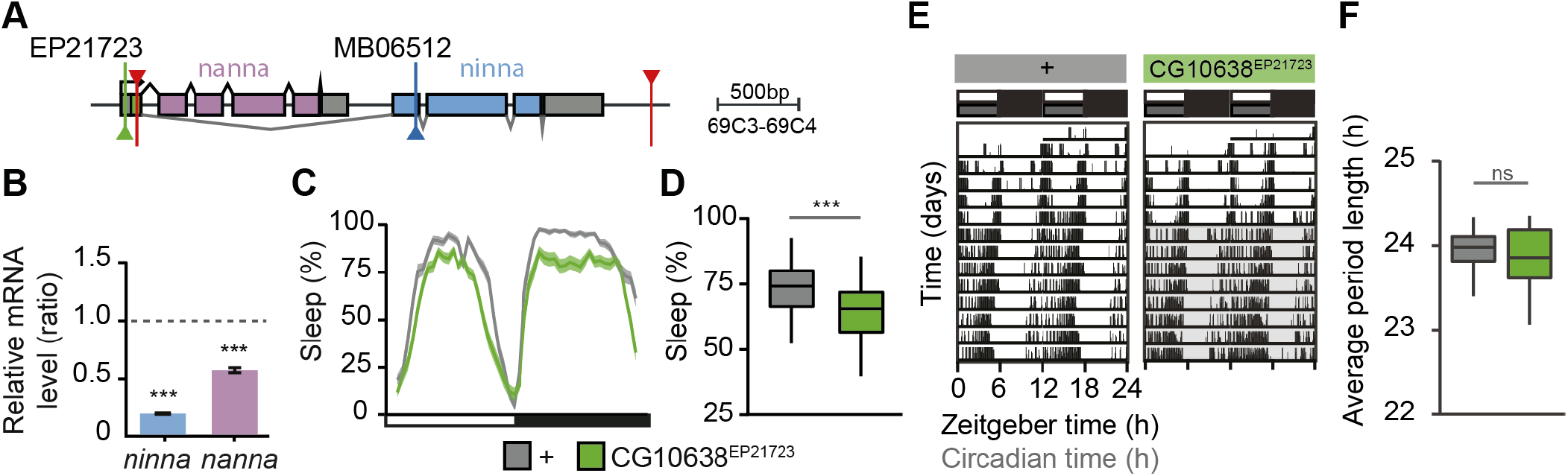
Disruption of the *ninna nanna* locus reduces homeostatic daytime sleep. **A**, Schema of the *ninna nanna* locus. Green arrow indicates P-element insertion of EP21723 mutant. **B**, mRNA levels for *ninna* (blue) and *nanna* (purple) transcripts in *ninnananna*^EP21723^ hypomorphic mutants, relative to expression in control flies (grey dashed line), n = 3 biological replicates per group (10 heads/sample), Student’s t-test. **C-D**, Sleep profile (**C**) and quantification (**D**) of P-element insertion mutant EP21723. WT control: n = 82, *ninnananna*^EP21723^: n = 90 flies, Wilcoxon signed rank test. **E-F**, Circadian rhythm of *ninnananna*^EP21723^ mutant flies. WT control: n = 15, *ninnananna*^EP21723^ mutants: n = 28 flies. **E**, Representative actograms of individual flies throughout the course of the experiment, respective genotypes indicated above the actogram. **F**, Quantification of free-running period length. Wilcoxon signed rank test. ns - not significant, *** P < 0.001. **B** Data represented as mean + SEM. **D**,**F**, Data are represented as median + IQR.

### Expression of Nanna tracks sleep pressure

Ninna nanna could regulate sleep by acting through process C or process S, or by affecting both processes (Fig. S2A). To test whether Ninna nanna tracks homeostatic and/or circadian sleep drive, we collected fly heads throughout the course of the 24 h cycle and assessed the expression of *ninna nanna* transcripts. If ninna nanna were involved in the circadian component of sleep regulation (process C), its recruitment would follow a circadian pattern similar to known clock genes such as period (Fig. S2B). Notably, the expression of the *nanna* transcript tracked regular sleep times (Fig. 3A), without, however, following a circadian pattern. *Nanna* expression was downregulated directly after sleep, both after night sleep (ZT0) and after “siesta” sleep (ZT9) – daytime sleep typically occurring around ZT6 – and upregulated at times coinciding with or immediately preceding regular sleep episodes, such as “siesta” sleep around ZT6 and night time sleep between ZT12 and ZT0 (Fig. 3A). This pattern almost faithfully reproduces changes in *bona fide* synaptic strength as previously quantified through the Bruchpilot synaptic protein (Fig. 3B) (17, 45–48) and suggests that expression of *nanna* encodes sleep pressure rather than circadian rhythm. We confirmed an increase of *nanna* transcript expression with prolonged time awake in male head extracts from flies collected immediately after sleep (ZT0) in comparison to flies collected after the waking day (ZT12, Fig. 3B), similar to expression levels of *bruchpilot*/*brp* (Fig. 3B) .

**Figure 3.**
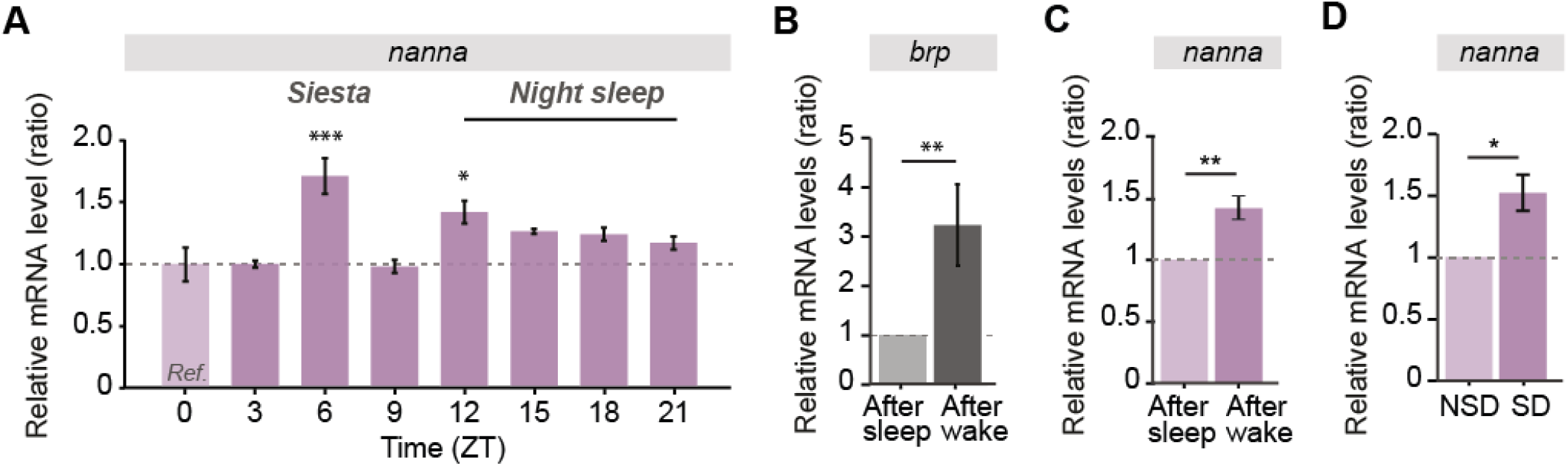
*Nanna* levels track sleep pressure. **A**, Transcript levels throughout the 24 h cycle, n = 3 per timepoint, ANOVA. mRNA levels during siesta (ZT6) differ from levels at neighboring timepoints (*post hoc* comparisons: ZT3:ZT6, ZT6:ZT9). mRNA levels in the beginning of night sleep (ZT12) differ from levels in the evening (*post hoc* comparison: ZT9:ZT12). **B-C**, Transcript levels at the end of the waking day (ZT12) and directly after sleep (ZT0). **B**, Transcript levels of *bruchpilot* (*brp*) after the waking day differ from levels directly after sleep. n = 3 per timepoint, Student’s t-test. **C**, Transcript levels of *nanna* after the waking day differ from levels directly after sleep. Sample size as in (**A**), Student’s t-test. **D**, mRNA levels of nanna transcripts. Heads were collected directly after overnight (12 h) sleep deprivation, control flies were kept under the same conditions but allowed to sleep and collected at the same time. n = 7 per treatment, Student’s t-test. **A-D**, One sample (n) constitutes one biological replicate (10 heads each). **B-C**, Heads were collected immediately after sleep (ZT0) or at the end of the waking day (ZT12). **A-D**, P < 0.05, ** P < 0.01, *** P < 0.001. Data represented as mean + SEM.

To test whether *nanna* expression responded to sleep pressure, we measured the expression of the *nanna* transcript immediately after overnight (12 h) sleep deprivation and found that *nanna* transcript levels increased in response to sleep deprivation in comparison to control flies that were allowed to sleep (Fig. 3C). We did not detect changes in the expression level of *ninna* or *period* transcripts in response to sleep deprivation (Fig. S2C,D). Together, these findings indicate that expression of *nanna* follows sleep pressure and suggest that nanna may be involved in the homeostatic regulation of sleep (process S).

### Nanna tracks homeostatic daytime sleep through the pair of inferior contralateral interneurons

To identify the substrate of nanna’s mechanism of action, we generated polyclonal antibodies raised against peptides replicating the non-conserved loop sequences (loop A and loop C in Fig. 1D), and stained adult brains to detect the expression pattern of each protein. Both proteins showed a very restricted staining in few, non-overlapping neurons across the brain. Anti-Nanna staining highlighted only a pair of symmetrical neurons located in the inferior lateral part of the brain — lateral to the subesophageal ganglion — and projecting contralaterally towards each other as well as towards the dorsal part of the protocerebrum (Fig. 4A). To better characterize the anatomy of *nanna*-expressing neurons, we screened the Vienna Tile GAL4 collection for driver lines highlighting a similar pattern, and identified the line VT026326 which specifically drives GAL4 in the two *Nanna*-positive neurons (Fig. 4B,C). As further confirmation that the VT026326 neurons express *nanna*, we knocked down *nanna* expression in those cells using RNAi and measured protein levels with anti-Nanna antibodies (Fig. 4D). Nanna protein levels assessed using the anti-Nanna antibody were reduced in flies with RNAi-mediated knock down of *nanna* in comparison to parental controls with intact *nanna* expression (Fig. 4E).

**Figure 4.**
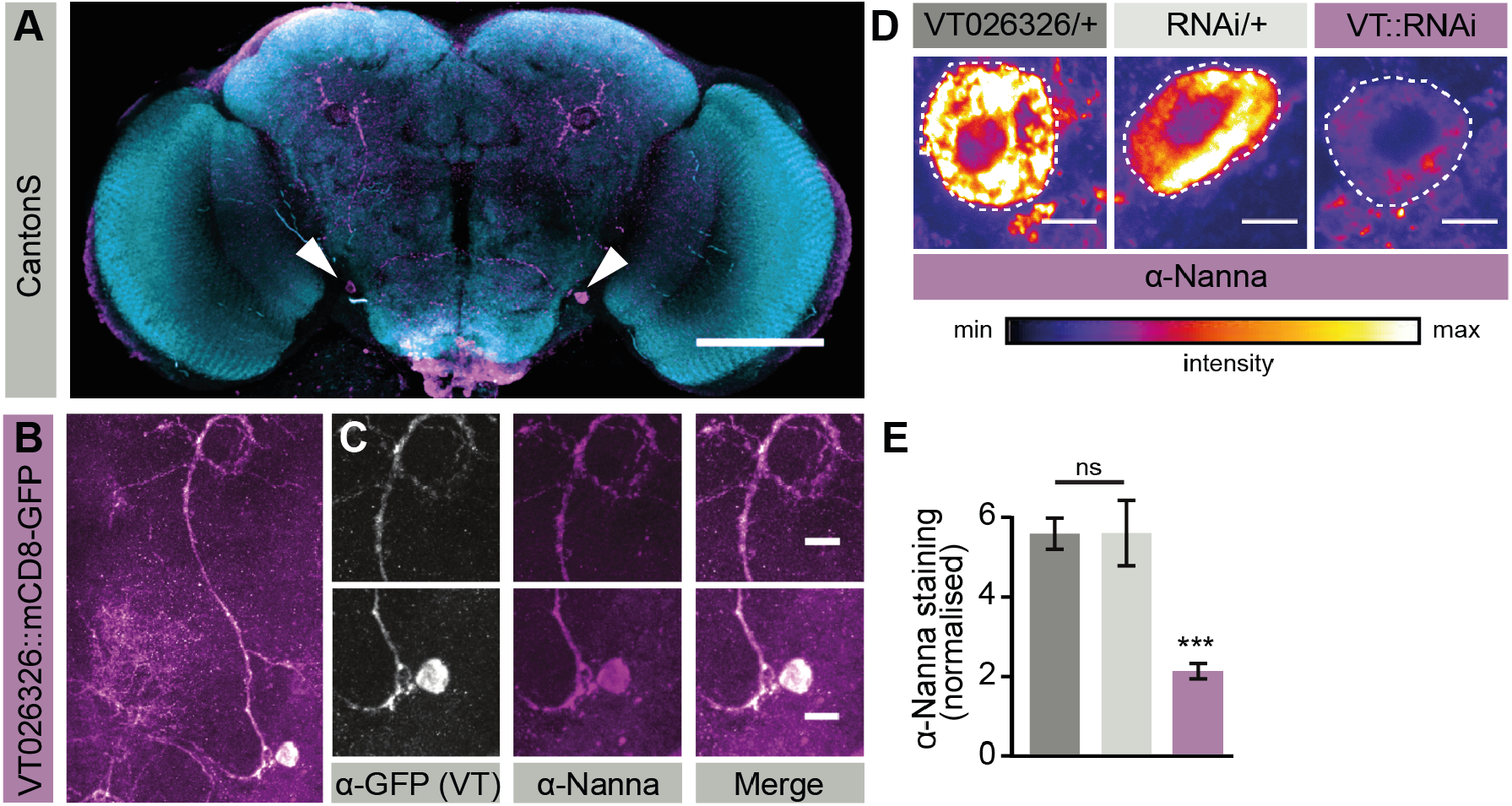
Nanna acts through the pair of inferior contralateral interneurons. **A**, Whole-brain anti-Nanna staining showing specific signal in a pair of symmetrical neurons (white arrows) projecting toward the middle protocerebrum. Blue: anti-nc82; purple: anti-Nanna. Scale bar 100 µm. **B-C**, mCD8-GFP driven by the VT026326-GAL4 driver (white) co-stained with anti-Nanna antibody (purple) with magnification of soma and neuronal processes (C). Scale bar 10 µm. **D**, Anti-Nanna staining of nanna neurons in RNAi knock down (purple header) or in parental controls (light and dark grey headers). Protein levels are visualized using the non-saturated Fiji fire palette. Scale bar 10 µm. **E**, Quantification of protein levels shown in the dashed area of (**D**), color code as in (**D**). VT026326-GAL4/+: n = 13, RNAi/+: n = 14, VT026326-GAL4::RNAi: n = 14 hemispheres. *** P < 0.001, Student’s t-test. Data represented as mean + SEM.

With their peculiar position and arborization (Fig. 4A), the *nanna*-expressing neurons are highly reminiscent of previously identified ventrolateral PDF-positive neurons (neuron 8 in (49)), i.e., a pair of interneurons known as inferior contralateral interneurons (ICLI) or inferior posterior slope neurons (IPS) that have been implicated in the regulation of sexual arousal and its effects on sleep homeostasis (50–52). To test their equivalence, we drove mCD8-GFP under control of the Tyramine receptor GAL4 (TyrR-GAL4, Fig. S3A) or under control of the myoinhibitory peptide GAL4 (MIP-GAL4 Fig. S3B) and co-stained with anti-Nanna antibodies. Indeed, *nanna*-expressing neurons were positive to both markers, confirming that they are IPS/ICLI neurons.

Interestingly, the anti-ninna antibody highlighted a different set of neurons, staining the PDF-expressing large lateral ventral neurons (l-LN_v_s) (Fig. S3C), a wake-promoting light-sensing cluster regulating nocturnal sleep in *Drosophila* (21, 53). We did not detect *nanna* expression in l-LN_v_s (Fig. S3D), and neither *ninna* nor *nanna* expression were detected in the other major group of PDF-expressing circadian neurons, the small lateral ventral neurons (s-LN_v_s) (lower panels in Fig. S3C,D). To confirm *ninna* expression in the l-LN_v_s, we used a GAL4-carrying Minos transposon (54) inserted in the first ninna-specific exon (MB06512, blue arrow in Fig. 2A) to drive the transmembrane mCD8-GFP protein as a membrane marker of *ninna* expression. Anti-GFP staining confirmed MB06512-GAL4 expression in the l-LN_v_s, showing a specific signal in the PDF-positive somata (Fig. S3E) as well as in elaborate medullar dendritic projections (Fig. S3F). Disruption of the ninna locus due to Minos transposon insertion in the MB06512 mutant did not produce a sleep phenotype (Fig. S3G,H). We did not detect expression of *ninna* in VT026326 neurons (not shown).

On the transcript level, *nanna* expression tracked sleep pressure (Fig. 3A,C,D). If ICLI neurons were the neural substrate of homeostatic sleep regulation by nanna, we hypothesized that nanna protein levels in ICLI neurons responded to sleep pressure. We conducted two sets of experiments to address this question: first, we examined whether Nanna protein levels in ICLI neurons changed as a function of sleep pressure. We compared antibody staining intensity between rested flies, at ZT0, and flies at the apex of their natural sleep drive, when sleep pressure is highest, at ZT12, and found that Nanna expression was indeed higher when sleep pressure was high (Fig. 5A,B). Secondly, we subjected wild-type flies to a 12 h regimen of sleep deprivation (SD) and compared Nanna levels between rested mock-treated flies and sleep deprived flies, collecting both at the same circadian time (ZT0). We observed a similar increase in Nanna protein levels in ICLI neurons (Fig. 5C,D).

**Figure 5.**
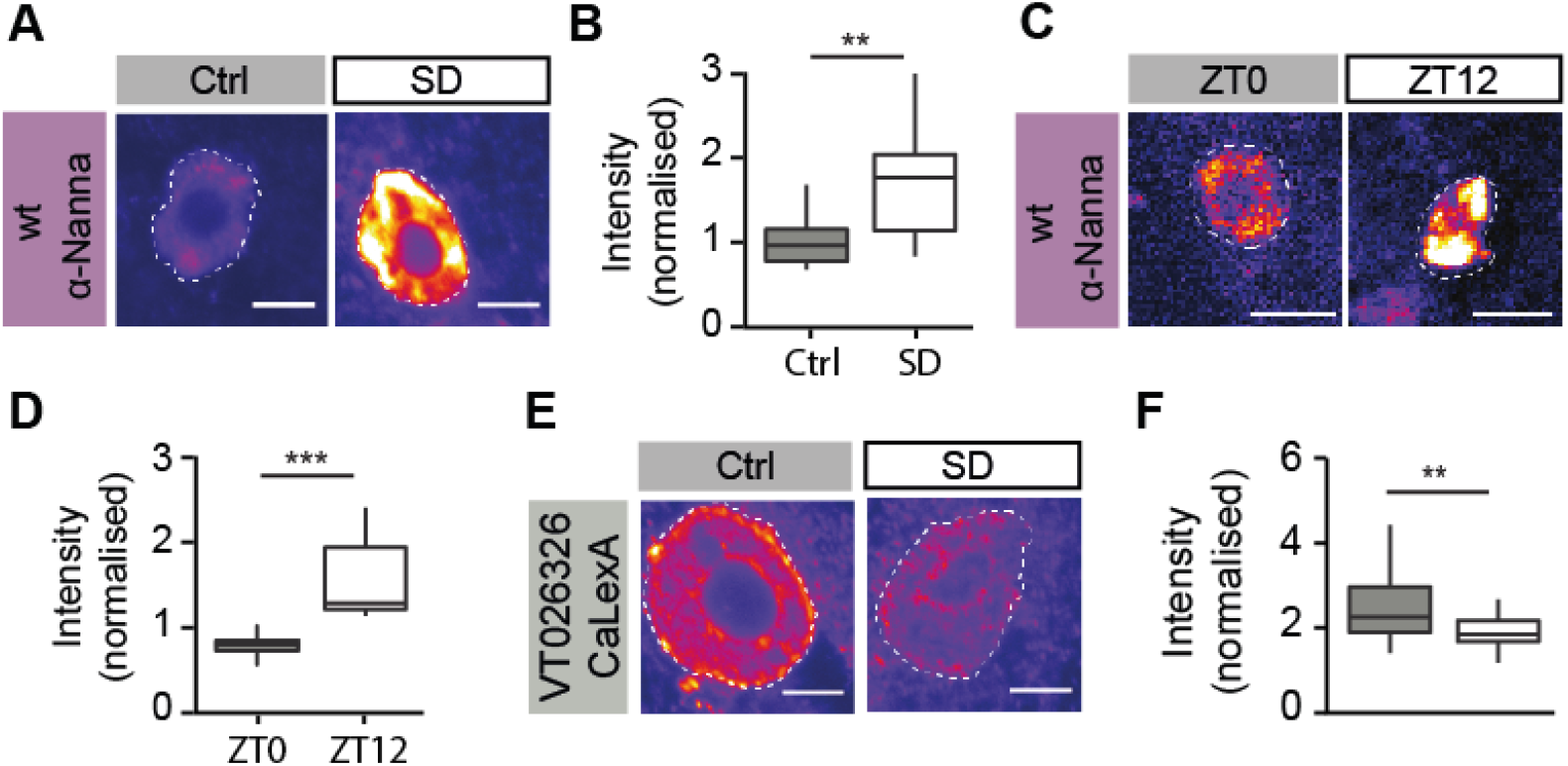
Nanna tracks the homeostatic sleep response of ICLI neurons. **A**, Anti-Nanna staining (fire palette) in wild-type CantonS flies at ZT0 and ZT12. Scale bar 10 µm. **B**, Quantification of protein staining in the dashed area highlighted in (**A**). ZT0: n = 8, ZT12: n = 9 hemispheres, Wilcoxon signed rank test. **C**, Anti-Nanna staining of rested flies (grey header) or flies that were sleep deprived for 12 h (white header), both groups collected at ZT0. Scale bar 10 µm. **D**, Quantification of the protein staining in the dashed area highlighted in (**D**). Ctrl: n = 14, SD: n = 15 hemispheres, Wilcoxon signed rank test. **E**, Representative image of anti-GFP immunostained flies expressing CaLexA in the nanna neurons using the VT026326 driver under control rested conditions (left, grey header) or after 12 h of sleep deprivation (right, white header). Scale bar 5 µm. **F**, Quantification of CaLexA signal in the dashed area of (**H**). Ctrl: n = 14, SD: n = 16 hemispheres, Wilcoxon signed rank test. ** P < 0.01, *** P < 0.001. Data represented as median ± IQR expressed.

To test whether an increase in Nanna protein levels corresponds to changes in the activity of ICLI neurons, we conducted two sets of experiments. In a first set of experiments, we employed the CaLexA system (55) to measure if sleep deprivation had any impact on the activity of *nanna*-expressing neurons. The CaLexA system relies on a fluorescent reporter gene whose transcription is spatially restricted by the GAL-UAS system and temporally responsive to intracellular calcium levels. Therefore, the intensity of detectable fluorescence in a genetically labelled neuron will proportionately reflect calcium influx and, ultimately, its firing rate (55). We expressed CaLexA in ICLI neurons using the VT026326-GAL4 driver, and compared the difference in fluorescence levels between rested control animals and animals subjected to 12 h of sleep deprivation. The fluorescence signal decreased in response to sleep deprivation (Fig. 5E,F), indicating that the activity of ICLI neurons decreases with sleep pressure, when nanna protein expression in ICLI neurons is conversely increased. These findings suggest that nanna may negatively regulate the activity of ICLI neurons at times of high sleep pressure to support sleep homeostasis.

### ICLI neurons are embedded in a sleep regulatory network

Light is the most important cue that entrains the circadian system of most animals, including fruit flies (56). Fruit flies are highly sensitive to light due to the circadian photopigment cryptochrome (CRY) which is widely expressed throughout the clock network. Major CRY- and PDF-expressing components of the clock network are the clusters of l-LN_v_ and s-LN_v_ neurons. The anatomical profile of ICLI neurons suggests that the light-sensitive input may derive from direct synaptic contact onto the s-LN_v_ cluster of PDF-expressing neurons in their dorsolateral input region (see Fig. 4A). As a first step in the characterisation of the circuit, we expressed mCD8-GFP under control of VT026326-GAL4 and concomitantly labelled the mushroom bodies by driving red fluorescent protein (RFP) under control of a lexA-specific driver (247-lexA) confirming that the *nanna*-expressing neurons project to and revolve around the peduncle of the mushroom bodies (Fig. 6A). The peduncle is heavily wrapped in glial sheaths that prevent synaptic contact with passing projections (57). Interestingly, the dorsolateral projection of ICLI neurons beyond the peduncle and towards the dorsolateral input region of the s-LN_v_ neurons stained positive for DenMark (Fig. 6B) and is hence dendritic in nature (58), suggesting that the *nanna*-expressing neurons receive input from this area. To test whether *nanna*-expressing neurons synaps directly onto s-LN_v_ neurons, we used the GRASP technique (GFP Reconstitution Across Synaptic Partners) (59) and expressed two split fragments of GFP under different promoters: a presynaptic fragment under control of a *pdf* driver (pdf-lexA) and a postsynaptic fragment under control of the VT026326-GAL4 line. We observed full GFP reconstitution, and consequent fluorescent signal, at the interface between the two neuronal populations in the dorsolateral protocerebrum (Fig. 6C,D). Together, these findings suggest that ICLI neurons receive direct synaptic input from PDF-expressing neurons, and most likely the s-LN_v_s. The s-LN_v_s are known to regulate circadian output and sleep through a small number of neuropeptides (60). Among these, the short neuropeptide F (sNPF) is believed to be the main sleep-promoting agent (61). To test the possible involvement of sNPF signalling in ICLI action, we co-stained adult brains with the anti-Nanna antibody and with an antibody directed against the sNPF receptor (anti-sNPFR). These stainings reveal that ICLI neurons indeed express the sNPF receptor and are likely to be receptive to sNPF signalling (Fig. 6E). In addition, ICLI neurons form axonal contralateral projections towards each other that lie above and penetrate into the subesophageal ganglion (SOG, Fig. 6F), indicating that ICLI neurons may output to neural clusters located in the SOG (Fig. 6G) – the seat of several neuronal clusters involved in homeostatic sleep regulation (23). Their input-output structure places ICLI neurons in an ideal position to link the light-sensitive, circadian sleep drive to the homeostatic sleep drive.

**Figure 6.**
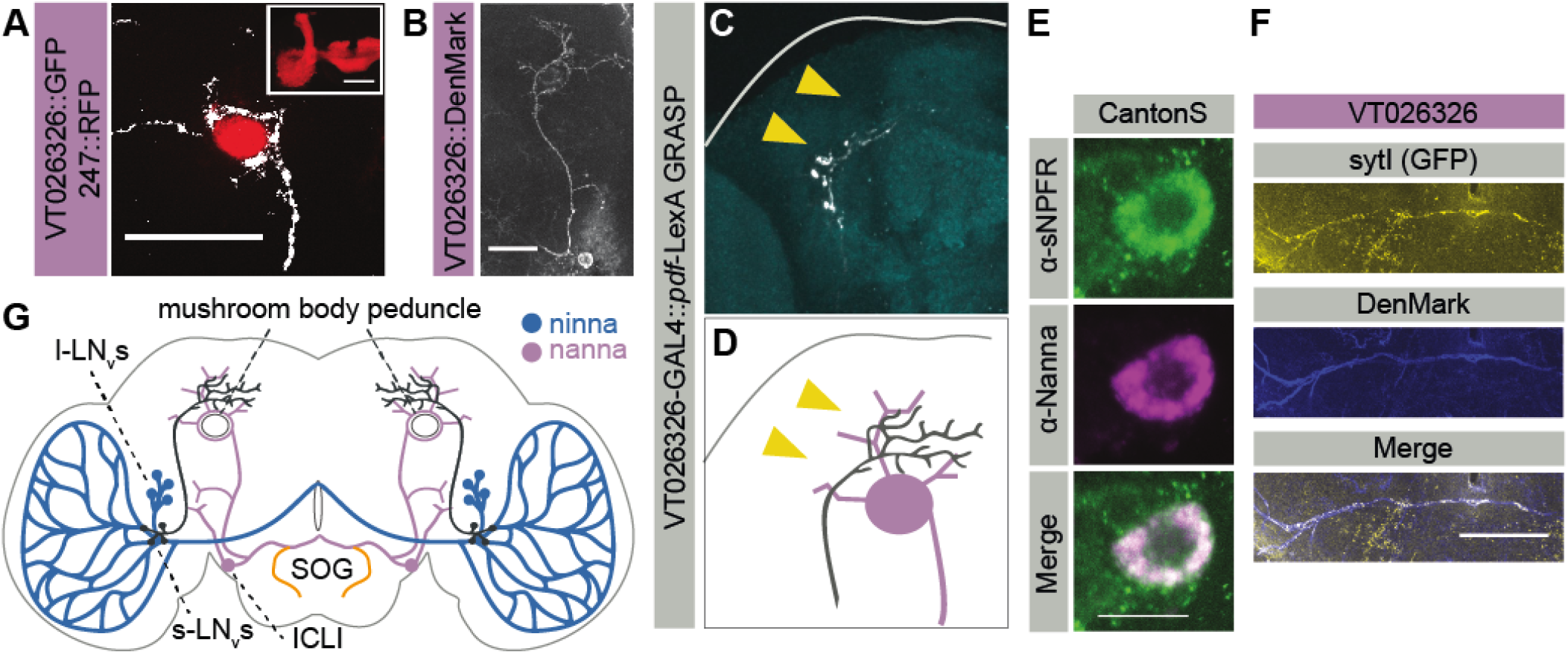
ICLI neurons are embedded in a sleep regulatory network. **A**, Co-staining of anti-RFP antibody (red) and anti-GFP antibody (white) in flies expressing mCD8-GFP under control of VT026326-GAL4 and RFP under control of the mushroom body specific driver 247-lexA. **B**, Expression of the dendrite specific UAS-DenMark (white) driven by the VT026326-GAL4 line along the dorsolateral projection of an ICLI neuron. **C-D**, GRASP signal revealing a widespread synaptic connection between pdf neurons and nanna neurons. **C**, Anti-GFP staining (white) of a whole-mount adult brain of pdf::spGFP110; VT026326::spGFP11 flies. The bottom panel shows the relevant area in (**D**), for spatial reference. Yellow arrows point to the relevant signal in both subpanels. **E**, Co-staining of anti-sNPFR (green) and anti-Nanna (purple) in wild type CantonS flies in the soma of an ICLI neuron. Scale bar 10 µm. **F**, Co-expression of the dendritic marker UAS-DenMark and the axonal marker UAS-sytI driven by VT026326-Gal4 along the contralateral projection of ICLI neurons in and above the SOG. Scale bar 20 µm. **G**, Schema depicting the expression of *ninna* and *nanna* in a coronal plane of the *Drosophila melanogaster* brain. *nanna* is expressed in ICLI neurons (purple) which project towards the dorsolateral protocerebrum wrapping around the mushroom body peduncle and also form axonal projections that cross over to the contralateral side (orange). ICLI neurons receive inputs from the s-LN_v_s (black) and send axonal projections (orange) into the subesophageal ganglion (SOG). **A-B**, Scale bar 20 µm.

## Discussion

Here, we characterised a novel gene, *ninna nanna*, that regulates homeostatic sleep in a transcript-specific manner. We identified a total of 12 AKR genes in *Drosophila melanogaster*, and found that only two were overexpressed in the brain: one brain-enriched AKR was Hyperkinetic, which serves as the *β* subunit of the voltage-gated potassium channel Shaker and regulates the activity level of dorsal fan-shaped body (dFB) neurons to support sleep homeostasis (39). The other brain-enriched AKR was *CG10638*, hereby named *ninna nanna*. Flies with disruption of the *ninna nanna* locus show reduced daytime sleep, a phenotype consistent with previous unbiased transcriptomic screenings which showed that *ninna nanna* expression correlates with reduced sleep in a genome-wide association study in *Drosophila* (62).

*ninna nanna* is differentially translated through a process of alternative splicing into two isoforms, Ninna and Nanna. According to transcriptomic screens, expression of *ninna nanna* was observed to oscillate with the clock (63, 64) and to be enriched in l-LN_v_s (65). Through newly generated antibodies, we confirm that Ninna is indeed expressed in the l-LN_v_s: a small group of PDF-positive circadian neurons previously described for their role in modulating nocturnal activity. The other transcript, *nanna*, is expressed in a pair of neurons that were previously identified as peptidergic neurons in *Drosophila* and in other species (49, 66, 67), termed intracontralateral interneurons (ICLI neurons). Importantly, we detected a higher expression of the nanna protein in ICLI neurons at times of high sleep pressure, concomitant with reduced neuronal activity in these neurons, suggesting that *nanna* expression either reflects or negatively influences the excitability of ICLI neurons. Future studies should address whether nanna acts in ICLI neurons to suppress their activeity and allow the animal to sleep and whether ninna plays a similar role in the circadian clock.

Most interestingly, *ninna*-expressing and *nanna*-expressing neurons connect into a unique functional circuit combining a light-sensing, circadian component to a component regulating sleep and other complex behaviour. To our knowledge, the *ninna nanna* gene is the first gene to be identified in which two alternatively spliced products are selectively expressed to form a functional neuronal circuit. Besides their striking complementary expression pattern, a second intriguing peculiarity of Ninna and Nanna is that they differ in their predicted use of co-factors. AKR proteins are extremely conserved in their tertiary structure and the co-factor binding pocket consists of a small number of amino acids with a well-described role (35, 68). Among these, two amino acids in the 8^th^ *β*-sheet were shown to be crucial for substrate selectivity: K260 and R268 (69, 70). Notably, Nanna bears substitutions in the two amino acids (K260 and R268) that are conserved amongst all eight analysed *Drosophila* species (Fig. 1D), strongly suggesting a conserved specificity for NADH over NADPH. Enzymes involved with NADH metabolism are reliable markers of wakefulness both in rodents (71) and in *Drosophila* (25), and levels of NADH correlate with neuronal activity (72, 73). A tantalising hypothesis is that the NAD^+^/NADH ratio may be a cellular marker for homeostatic sleep pressure, in a conceptually similar way to what has been proposed for adenosine (74), and Nanna may specifically act as a redox sensor for NADH in ICLI neurons to scale neural activity to homeostatic sleep pressure signalled by the NAD^+^/NADH ratio. A converse mechanism for NADP^+^/NADPH ratio has been proposed for another AKR, Hyperkinetic (39). Notably, both *hyperkinetic* and *ninna nanna* produce only a moderate sleep phenotype when perturbed ((29, 39) and this study), conspicuously less dramatic to that observed with disruption of *shaker*/*minisleep* (25) or its other regulator, *sleepless* (27, 28). These observations indicate either redundancy with other sensors or that the role of oxidative stress plays only a limited part in homeostatic regulation. Comparative analysis showed that Nanna does not belong to the K_v*β*_ family of AKRs, and we could not find any phenotypical epistasis between *shaker* or *hyper-kinetic* and *nanna* – neither in terms of sleep levels nor in terms of leg shaking (data not shown) - indicating that the cellular sleep-regulatory functions of Nanna are distinct from those of Hyperkinetic.

Expression data suggest that Ninna, on the other hand, may act as NADP^+^/NADPH sensor in the circadian clock. In recent years, the NADP^+^/NADPH ratio has been implicated in the regulation of circadian clocks in various models: in enucleated red blood cells, redox cycles have been shown to set circadian rhythms in a transcription-independent manner (75); in the main mammalian pacemaker neurons (the SCN) NADP(H) and NAD(H) were shown to cycle and reinforce the robustness of the molecular clock by ultimately regulating the activity of voltage-gated K^+^ channels (76, 77). In mammals, NAD(P)(H) regulates the binding between, and hence the activity of, NPAS2 and BmalI, two key regulators of the molecular clock (78, 79). NPAS2, in particular, is directly modulated by binding with NADPH and plays an important role not just in regulating the clock, but also in sleep regulation (44, 80).

Taken together, the results presented here depict a possible archetypal circuit for sleep regulation, integrating a circadian drive (process C) and a homeostatic drive (process S). The circuit is composed of two main sensors: the circadian sensor uses an external input (light) and internal reinforcers (NADP^+^/NADH ratio) to gauge circadian information and modulate the output of s-LN_v_s to the second sensor, a homeostatic somnostat expressed in ICLI neurons. This latter segment of the circuit integrates the internal information regarding sleep drive with the internal information regarding arousal and ultimately drives the animal towards sleep or activity. This *ninna nanna* circuit is unlikely to be the main sleep regulatory circuit in *Drosophila*. Its description rather speaks for a scenario in which sleep is regulated by a network of smaller circuits, integrating circadian drive and internal drive. For the brain – and the body – the major advantage of using a molecular marker of sleep pressure, such as NAD(H), is that a global signal could be used to inform the entire organism of the current sleep homeostatic drive. This has been a strength of the adenosine hypothesis so far and our results suggest a possible functional analogue.

## Author contribution

AP performed all experiments and led the project; GFG designed and supervised the project, and obtained the funding. Both authors wrote the manuscript.

## Acknowledgement

The research leading to these results has received funding from BBSRC through BB/M003930/1. AP was supported by a PhD scholarship of the Edmond J. Safra foundation. We thank the Facility for Imaging by Light Microscopy (FILM) at Imperial College London, part-supported by funding from the Wellcome Trust (grant 104931/Z/14/Z) and BBSRC (grant BB/L015129/1). We would like to thank Ralf Stanewsky, Marc Dionne, Scott Waddell, Kweon Yu, and Jongkyeong Chung for generously sharing reagents with us; Jess Sharrock, Solweig Schlumberger, Ho Yan Grace Wu, Clara Castello, and Kallista Chan for their technical support; Esteban Beckwith and Erhard Hohenester for comments on the manuscript. Special thanks to all the members of the Gilestro, Southall and Dionne laboratories – particularly Katrin Kierdorf - for their constant and invaluable input and support.

## Materials and Methods

### Experimental model and subject details

Flies were raised under a 12 h light:12 h dark (LD) regimen at 25^*°*^C on standard corn and yeast media. The following lines were used in the study: CantonS and pdf-lexA from Dr. Ralf Stanewsky (Muenster University, Germany); w1118 from Vienna Drosophila RNAi centre (VDRC, Austria); elav-GAL4, pdf-GAL4, TyrR-Gal4, UAS-DenMark, ninna nan-naEP21723, UAS-GFP1-10 and UAS-GFP11 from Bloomington Drosophila Stock Centre (BDSC, Indiana, USA); 247-lexA and lexAop-RFP from Scott Waddell (CNCB, UK); MIP-Gal4 from Jongkyeong Chung (Seoul National University); UAS-CaLexA from Marc Dionne (ICL, UK); all the UAS-RNAi lines and the VT026326-GAL4 line from VDRC.

### Bioinformatics of Ninna Nanna structure

The tertiary structure of Ninna and Nanna was inferred by submitting the respective protein sequences to Phyre2 (81). For Ninna, the analysis returned a structure modelled with 100 % confidence upon the crystal for the human aldose reductase (PDB 1US0, primary sequence identity 46 %). For Nanna, the analysis returned a structure modelled with 100 % confidence upon the crystal for the aldose reductase of Schistosoma Japonicum (PDB 4HBK, primary sequence identity 42 %). The sequences of eight Drosophila species shown in Figure S1 were obtained through flybase, automatically aligned through ClustalW, and manually curated.

### Sleep recording

The activity of individual flies was recorded using Drosophila Activity Monitors (DAM2, TriKinetics, Waltham, MA, USA) and acquired by the DAMSystem3 Data Acquisition Software supplied by the manufacturer. For regular baseline recordings, at least 6 days were recorded, whereby the first two days were regarded as the habituation period and discarded for analysis.

### Antibodies and immunofluorescence

Polyclonal antibodies against Ninna and Nanna were raised in guinea pigs against synthetic peptides. Two peptides were generated for each isoform, corresponding to the exposed loop regions as identified from the Phyre2 model. These are: DDNTLLP-KNEDDVLQ (ab 1), YTPLGKPKPDIQKPD (ab 2), EDLM-PHENGQLRTND (ab 3), GRPKACNPLPDYYTD (ab 4). Peptide antibodies were synthetised and purified by Eurogentec (anti-peptide programme AS-DOUB-LXPGUI). Anti-PDF (PDF C7), anti-BRP (nc82) are from the *Drosophila* hybridoma Bank, DSHB. Anti-sNPFR antibody is a gift from K. Yu (KRIBB, S.Korea). Anti-GFP and anti-RFP are from abcam (ab290 and ab62341 respectively). All secondary antibodies are from abcam. Brains were stained and fixed as previously described (45).

### CaLexA measurements

Animals were grown and treated in the same conditions as in behavioural experiments. After mechanical sleep deprivation, animals were anaesthetised and their brains were dissected and fixed as previously described (45). For CaLexA measurements, fly brains were immunostained with anti-GFP (1:400, ab290 Abcam). Images were taken under 40 times magnification and analysed in Fiji/ImageJ (82). To measure signal intensities, a maximal intensity projection of all the stacks. Intensity on an adjacent non-labelled region was measured and subtracted.

### Sleep deprivation

Mechanical sleep deprivation was conducted by placing the flies on top of a laboratory shaker controlled by an Arduino timer activated in pulses of 5 to 30 seconds at pseudo-random intervals of 1 to 7 minutes (Arduino code and instructions on https://github.com/gilestrolab/fly-sleepdeprivator).

### Quantification and statistical analysis

All data analysis was performed in R. Behavioural data were analysed with the R package Rethomics (https://github.com/gilestrolab/rethomics). Datasets were tested for normal distribution using Shapiro-Wilk normality test at an *α* level of 0.05. For normally distributed data sets, parametric tests were used to test for differences between conditions. These include either Welsh two sample t-test for paired independent samples, or ANOVA for comparisons across multiple groups and for co-variance analysis. To visualise normally distributed, parametric data sets, results were represented in bar plots. Throughout, bar height reflects the average and error bars reflect the standard error of the mean (“SEM”). For non-normally distributed datasets, non-parametric tests were used to test for differences between conditions. These include either Wilcoxon signed-rank test for paired independent samples, or Kruskal-Wallis test for comparisons across multiple groups and for co-variance analysis followed by Dunn’s post-hoc test. To visualise non-normally distributed, parametric data sets, results were represented in box plots.

For ethograms, bootstrap re-sampling with 5000 replicates, was performed in order to generate 95 % confidence interval (83) (shadowed ribbons around the mean in the figures). All experiments were replicated two to five times. In all figures, Ns represent the total number of flies over all experiments. Statistics were done on aggregated data. Outliers were never excluded. Flies that died during the course of the experiment were excluded from all analysis. All figures were generated in R (*ggplot2* package). For all boxplots, the bottom and top of the box (hinges) show the first and third quartiles, respectively. The horizontal line inside the box reflects the second quartile (median). Tuckey’s rule (the default), was used to draw the “whiskers” (vertical lines): the whiskers extend to last extreme values within ± 1.5 IQR, from the hinges, where IQR is Q3-Q1.

## Supplementary figures

**Figure S1.**
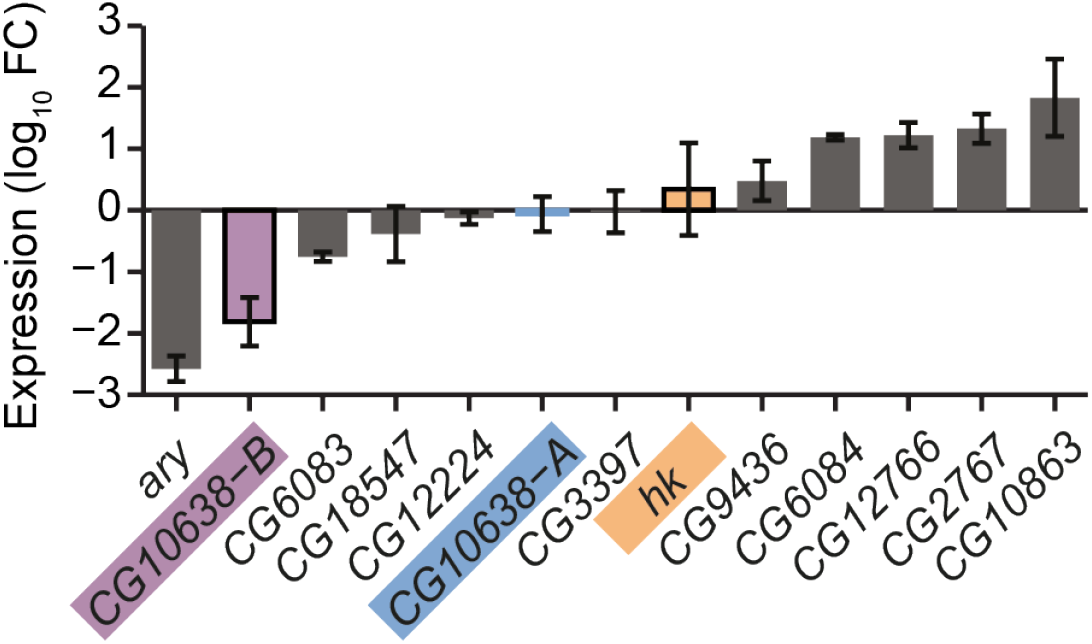
The AKR genes in *Drosophila* melanogaster -extended. Related to Figure 1. Expression of AKRs in the head of the fly. n = 3-5 independent biological samples consisting of 10 heads each. Data represented as mean + SEM.

**Figure S2.**
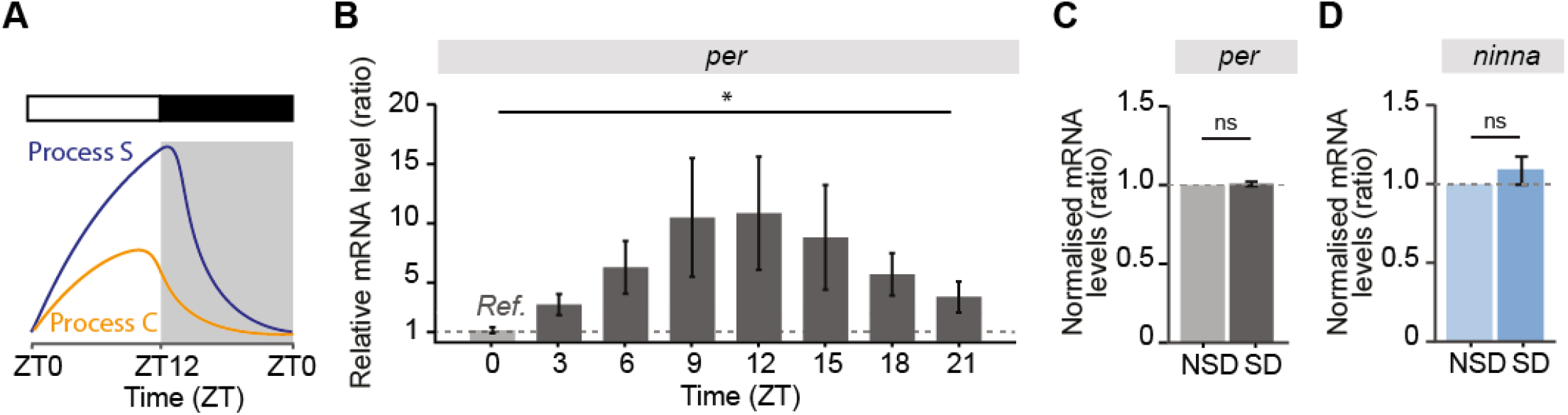
Expression of ninna and per transcripts. Related to Figure 3. **A**, Schema of circadian (process C) and homeostatic (process S) sleep regulation. **B**, mRNA levels of *period* (*per* ) transcripts throughout the 24 h cycle, n = 3 per timepoint. * P < 0.05, ANOVA. **C**, mRNA levels of *per* transcripts, n = 7 per treatment, ns – not significant, Student’s t-test. **D**, mRNA levels of *ninna* transcripts, n = 7 per timepoint, ns – not significant, Student’s t-test. **B-D**, One sample (N) constitutes one biological replicate (10 heads each). Data represented as mean + SEM.

**Figure S3.**
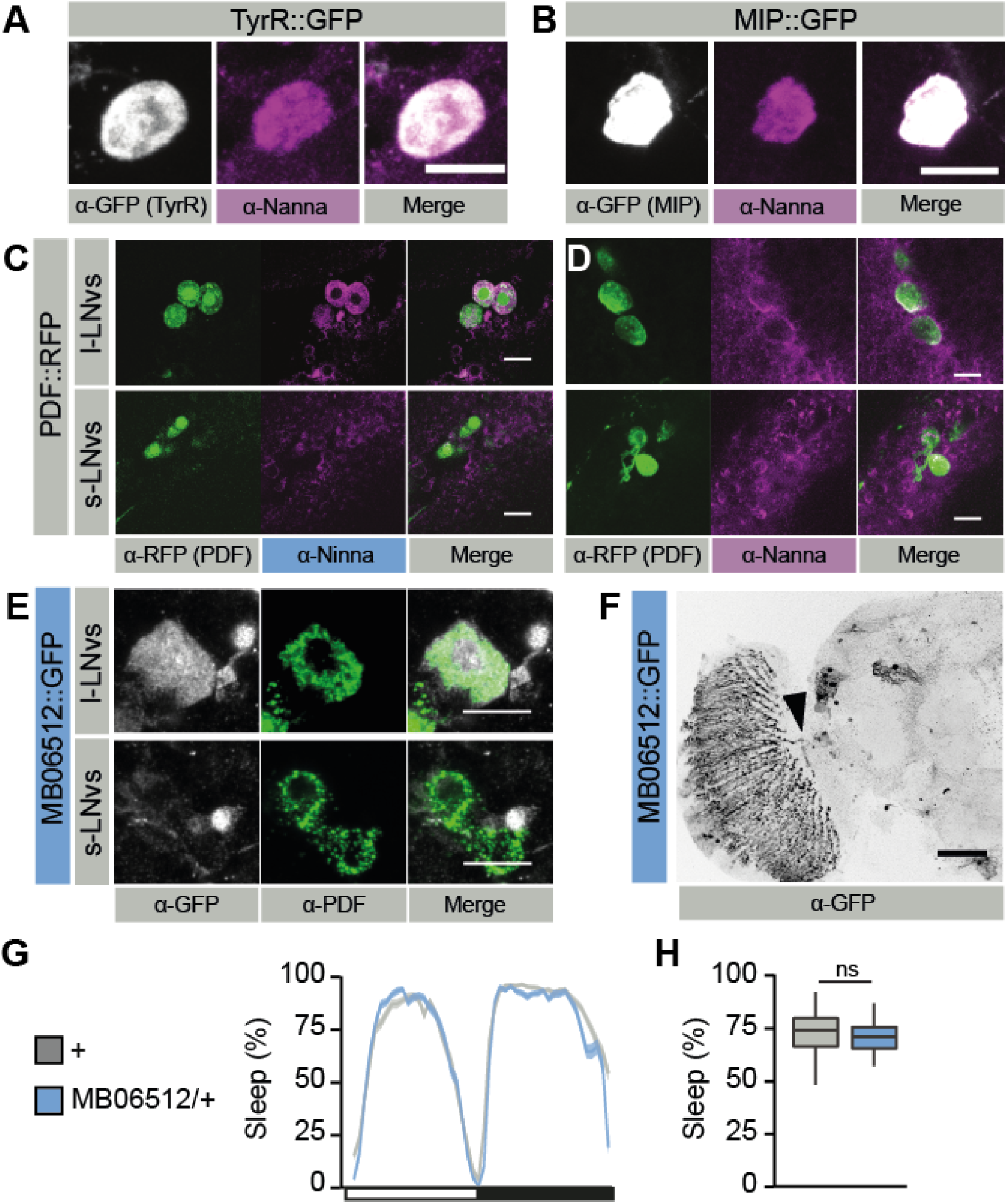
Additional behavioral assessments and antibody stainings related to Figures 4 and 5. **A-B**, Co-staining of anti-Nanna (purple) and anti-GFP (white) in flies expressing mCD8-GFP under control of the TyrR-GAL4 driver (**A**) or of the MIP-GAL4 driver (**B**). **C-D**, Expression of ninna in circadian pacemaker neurons. **C**, Ninna is expressed in l-LNvs but not s-LNvs. Flies expressing RFP under control of the pdf-GAL4 driver were stained with anti-Ninna antibody (purple) and anti-RFP antibody (green). **D**, Same as (**C**), but with anti-Nanna antibody. **E**, Flies expressing mCD8-GFP under control of the MiMic transposon MB06512, inserted in the ninna locus, co-stained using anti-GFP (white) and anti-PDF (green) antibodies. Ninna staining is visible in the l-LNvs. **F**, mCD8-GFP driven by MB06512 (black) shows expression in the l-LNvs soma (black arrow) and the elaborated projection in the medulla. Scale bar 40 µm. **G-H**, Sleep of MB06512 with sleep profile (**G**) and quantification (**H**). WT control, n = 63, MB06512/+, n = 82 flies. ns – not significant, Wilcoxon signed-rank test. Data represented as median + IQR. **A-D**,**E**, Scale bar 10 µm.

